# Fibroblast growth factor receptors function redundantly during zebrafish embryonic development

**DOI:** 10.1101/501023

**Authors:** Dena M. Leerberg, Rachel E. Hopton, Bruce W. Draper

**Affiliations:** Department of Molecular and Cellular Biology, University of California, Davis Davis, California, United States of America; Present address: Section of Cell and Developmental Biology, University of California, San Diego, La Jolla, California, United States of America; Present address: Institute of Molecular Biology, University of Oregon, Eugene, Oregon, United States of America

**Keywords:** Fibroblast growth factor signaling, posterior mesoderm, pectoral fin, midbrain-hindbrain boundary, viscerocranium, neurocranium

## Abstract

Fibroblast growth factor (Fgf) signaling regulates many processes during development. In many cases, one tissue layer secretes an Fgf ligand that binds and activates an Fgf receptor (Fgfr) expressed by a neighboring tissue. Although many Fgf ligands have known requirements in development, less is known about the requirements for the receptors. We have generated null mutations in each of the five *fgfr* genes in zebrafish. Considering the many requirements for Fgf signaling throughout development and that null mutations in the mouse *Fgfr1* and *Fgfr2* genes are embryonic lethal, it was surprising that all zebrafish homozygous mutants are viable and fertile, with no discernable embryonic defect. Instead, we have discovered surprising complexity of the Fgf pathway, where multiple receptors are involved in coordinating developmental processes. For example, mutations in the ligand *fgf8a* cause loss of the midbrain-hindbrain boundary, whereas in the *fgfr* mutants, this phenotype is only seen in embryos that are triple mutant for *fgfr1a;fgfr1b;fgfr2*, but not in any single and double mutant combinations. We show that this apparent *fgfr* redundancy is also seen during the development of several other tissues, including posterior mesoderm, pectoral fins, viscerocranium, and neurocranium. These data therefore begin to define the Fgfrs that function with a particular Fgf ligand to regulate early development in zebrafish.

**Summary statement:** Analysis of fibroblast growth factor receptor mutants in zebrafish shows that functional redundancy assures robustness of a core developmental signaling pathway.

## Introduction

The coordination of cellular events that drive developmental processes requires robust communication between cells. One common communication mechanism is the Fibroblast growth factor (Fgf) signaling pathway, which involves a diffusible Fgf ligand that is secreted into the extracellular space where it interacts with heparan sulfate and an Fgf receptor (Fgfr) (reviewed in (Mohammadi et al., 2005)). Fgfrs are single-pass transmembrane proteins composed of an extracellular region containing immunoglobulin (Ig) domains and an intracellular tyrosine kinase domain. Upon ligand binding, receptor dimerization leads to transphosphorylation-dependent activation of the kinase domain (Lemmon and Schlessinger, 1994). Fgf signaling can activate several intracellular signal transduction pathways, including the MAPK, PLCγ, and PI3K/Akt cascades (Pawson, 1995). While Fgf signaling can result in transcriptional changes, the ultimate cellular response depends on context and ranges from proliferation to migration and differentiation (reviewed in (Powers et al., 2000)).

Although the Fgf signaling pathway likely arose in eumetazoans (Bertrand et al., 2014), it has become quite elaborate in more complex animals. For example, the mammalian genome contains 22 ligand and 4 receptor genes, whereas the zebrafish genome has 31 ligand and 5 receptor genes (Ornitz and Itoh, 2001). These genes are expressed widely throughout both developing and mature tissues, often in overlapping domains (Ota et al., 2010). Tissue culture-based experiments have indicated that individual Fgf ligands have some degree of preference for the receptors they activate. This preference seems to be conferred by interactions between the glycine box, a ∼10 AA stretch near the C-terminus of an Fgf ligand (Luo et al., 1998) and the third Ig domain (IgIII) of the receptors. Interestingly, FGFR1, FGFR2, and FGFR3 in mammals and Fgfr1a and Fgfr2 in zebrafish are subject to alternative splicing in this IgIII domain, and these alternative isoforms have different affinities for particular ligands (Chellaiah et al., 1994; Johnson et al., 1991; Werner et al., 1992; Yeh et al., 2003).

Previously, studies have investigated the roles of Fgf signaling by disrupting the function of particular ligands. In zebrafish alone, the function of many Fgf ligands during early development has been determined using genetic mutation or morpholino knockdown. In some cases, disrupting a single Fgf gene leads to a developmental defect. For example, disrupting *fgf24, fgf10a*, or *fgf16* signaling results in the absence of pectoral fins, whereas loss of *fgf8a* leads to midbrain-hindbrain boundary (MHB) defects (Brand et al., 1996; Fischer et al., 2003; Manfroid et al., 2007; Nomura et al., 2006; Reifers et al., 1998). In other contexts, however, Fgf ligands appear to function redundantly. While both *fgf8a* and *fgf24* single mutants have normal mesoderm development, disrupting both ligands simultaneously leads to loss of posterior mesodermal derivatives and a consequent shortening of the embryonic axis (Draper et al., 2003). Similarly, simultaneous loss of both *fgf8a* and *fgf3* leads to severe defects in pharyngeal pouch development, whereas this tissue develops normally in either single mutant. These and similar data suggest that genetic redundancy in the Fgf signaling components creates a robust developmental system (Brand et al., 1996; Draper et al., 2003; Fischer et al., 2003; Reifers et al., 1998).

In contrast to the known requirements of many Fgf ligands during development, comparatively less is known about the requirements for specific Fgfrs. In the mouse, null mutation of *Fgfr1* or *Fgfr2* is embryonic lethal (Arman et al., 1998; Deng et al., 1994; Yamaguchi et al., 1994), whereas tissue-specific disruption of these genes reveals their roles during later development in limbs and/or brain (Trokovic et al., 2003; Xu et al., 1999a; Xu et al., 1998). By contrast, *Fgfr3* and *Fgfr4* homozygous mutants are embryonic viable, though *Fgfr3* single mutants have skeletal dysplasia, and *Fgfr3*;*Fgfr4* double mutants have defective lung development. These latter results suggest that, like certain Fgf ligands, in some developmental contexts Fgf receptors also function redundantly (Colvin et al., 1996; Deng et al., 1996; Weinstein et al., 1998). The zebrafish genome contains single copies of *fgfr2-4* orthologs, and two copies of an *fgfr1* ortholog, called *fgfr1a* and *fgfr1b*, that appear to have arisen during the teleost-specific whole genome duplication (Rohner et al., 2009). In contrast to several *fgf* ligand mutants, which have been isolated in phenotype-based forward genetic screens for recessive mutations, only one mutation in an Fgf receptor has been identified in a recessive screen, a result that could be explained if Fgf receptors function redundantly. The one exception is *fgfr1a*, where a point mutation in the kinase domain, proposed to be a strong hypomorph, is embryonic viable but has defects in scale formation during juvenile development.

To determine the precise requirements of each Fgf receptor during embryonic development, we have produced loss-of-function mutations in each of the five zebrafish *fgf* receptor genes. Because of the known requirements for Fgf signaling during early zebrafish development, we expected that some of the receptor mutants would phenocopy known Fgf ligand mutants. However, we found that all single mutants are viable with no overt embryonic phenotypes; instead we discovered that only certain double and triple mutant combinations have developmental defects in the posterior mesoderm, brain, pectoral fin, and pharyngeal arch derived cartilages. These findings suggest significant genetic redundancy between various Fgf receptors, and indicate that some ligands are capable of activating signaling through as many as three different receptors.

## Results and Discussion

### Generation of mutant alleles

To determine the function of Fgf receptors in zebrafish development, we generated a null allele for each gene using the CRISPR/Cas9 gene editing system. Fgf receptors are composed of an extracellular ligand-binding domain, a single transmembrane domain, and an intracellular kinase domain. We used single-stranded guide RNAs to target Cas9 endonuclease to the 5’ end of each gene to induce frameshift-causing indel mutations, which were confirmed by sequencing genomic DNA and then cDNA to assess the possibility of exon skipping that could result in a truncated, but functional, protein ((Mou et al., 2017; Sharpe and Cooper, 2017); Table 1; Fig. 1A)). A 127-bp insertion into *fgfr1a* exon 5 was the only mutation that resulted in exon skipping. However, because exon 5 is not a multiple of three (173 bp), the misspliced transcript goes out of frame at the aberrant splice junction. The predicted peptides that can be translated from each of the five mutant alleles are illustrated in Fig. 1A.

**Table 1.** Nature of CRISPR/Cas9-generated alleles

| Gene | Allele | Nature of genomic disruption | Exons skipped during splicing | WT peptide length (AA) | Predicted length of resulting peptide (AA) | # of AA in frame |
| --- | --- | --- | --- | --- | --- | --- |
| *fgfr1a* | uc61 | 127 bp inserted into exon 5 | Exon 5 | 810 | 202 | 138 |
| *fgfr1b* | uc62 | 5 bp deleted from exon 6 | - | 741 | 220 | 220 |
| *fgfr2* | uc64 | 47 bp deleted from exon 3 | - | 838 | 76 | 71 |
| *fgfr3* | uc51 | 50 bp deleted from exon 3 | - | 821 | 95 | 49 |
| *fgfr4* | uc50 | 4 bp deleted from exon 3 | - | 922 | 134 | 64 |

**Fig. 1.**
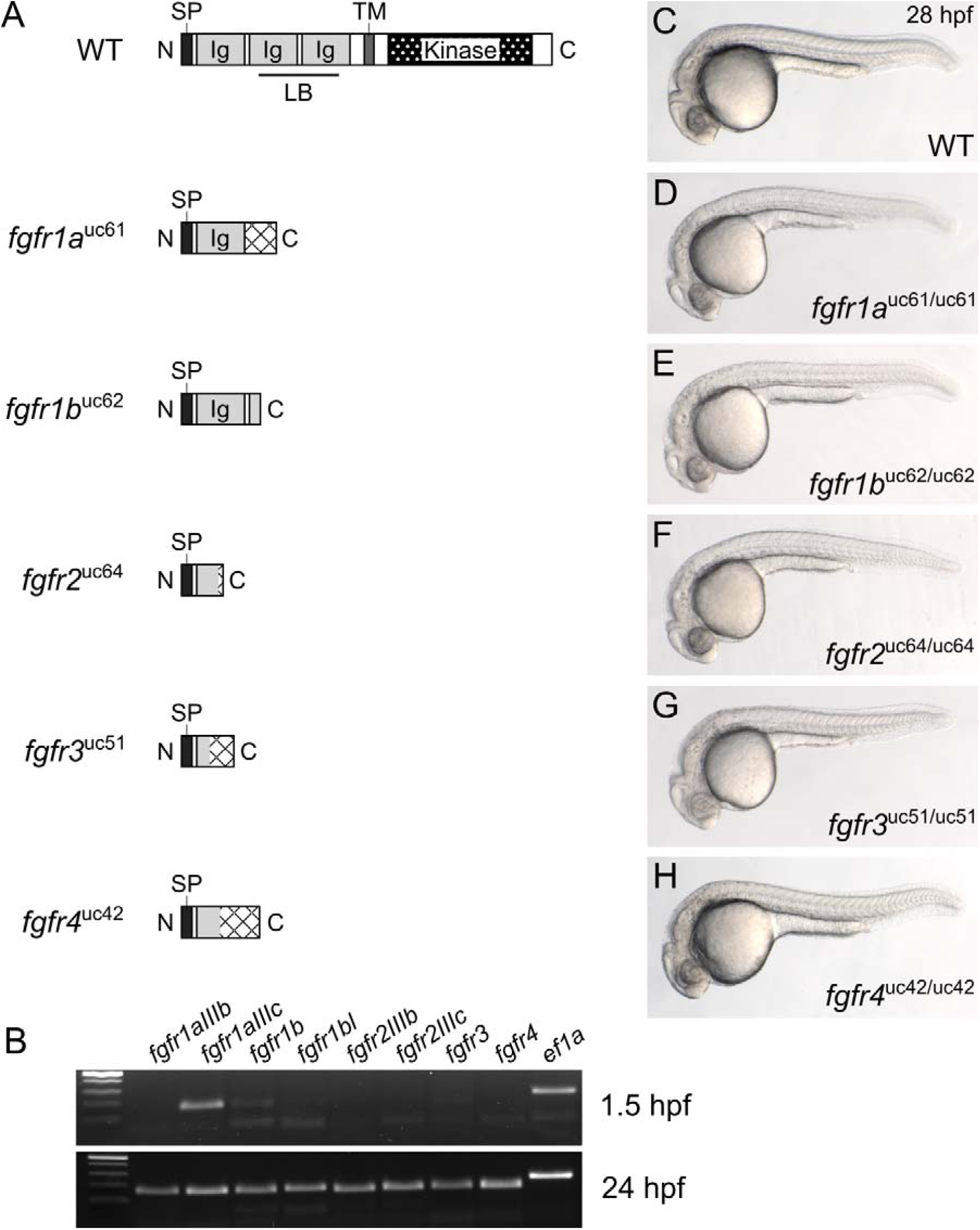
Fgf receptor mutants are embryonic viable. (A) RT-PCR of wild-type embryos. While all *fgfr* isoforms are detected in 24 hours post fertilization (hpf) embryos (post-zygotic genome activation, bottom panel), only *fgfr1a* (isoform IIIc) and *fgfr1b* are detected in 1.5 hpf embryos (pre-zygotic genome activation, top panel). *ef1a* is shown as a positive control. (B) Schematic diagram of a typical full-length Fgfr protein and the predicted truncated peptides resulting from the *fgfr1a*^*uc*61^, *fgfr1b*^*uc*62^, *fgfr2*^*uc*64^, *fgfr3*^*uc*51^, and *fgfr4*^*uc*42^ alleles. SP = signal peptide; LB = ligand-binding domain; Ig = immunoglobulin; TM = transmembrane domain; hatching indicates missense amino acids. (C-H) Lateral view of ∼28 hpf wild-type (WT, C) or homozygous mutant (D-H) embryos. Anterior is to the left, dorsal is up.

Fgf signaling is involved in many developmental processes, and all five *fgfr* genes are expressed during zebrafish embryogenesis (Fig. 1B; (Ota et al., 2010)). It was therefore surprising that all five *fgfr* homozygous mutants were embryonic viable and had no apparent defects when compared to wild-type siblings (Fig. 1C-G). Because there is precedent in zebrafish for zygotic defects being masked by maternally provided gene products (e.g. (Giraldez et al., 2005)), we asked if any of the Fgf receptor mRNAs are maternally provided and found that indeed *fgfr1a* RNA (specifically the splice isoform *IIIc*), and to a lesser extent, *fgfr1b* RNA, are maternally provided (Fig. 1B; (Ota et al., 2010; Rohner et al., 2009; Scholpp et al., 2004)). It was therefore possible that this maternal contribution could explain the absence of a phenotype when only the zygotic *fgfr* expression is lost. To test this, we produced maternal-zygotic mutants (MZ), which are mutant embryos derived from homozygous mutant mothers and therefore lack both maternal (M) and zygotic (Z) gene products. We found that MZ*fgfr1a* and MZ*fgfr1b* single mutant embryos were phenotypically wild-type (Fig. S1). Thus, the absence of phenotypes for any of the receptor single mutants is not likely due to rescue by maternally provided gene products. These data also suggest that the receptor genes act in a redundant or compensatory manner during early development.

Previously, a point mutation in *fgfr1a*, called *fgfr1a(t3R05H*), was shown to affect juvenile scale development in zebrafish (Rohner et al., 2009). Animals homozygous for this mutation, which affects a conserved arginine in the intracellular kinase domain and is predicted to be a strong hypomorph, develop with fewer flank scales, and the remaining scales are significantly larger than those of wild-type animals. We therefore asked if a similar scale phenotype in present in animals homozygous for *fgfr1a(uc61)* null mutations. Indeed, three of five *fgfr1a(uc61)* mutants analyzed had noticeably larger flank scales compared to wild-type siblings (dotted outlines, Fig. S2). However, we did not observe the severe reduction in scale number reported for *fgfr1a(t3R05H*) mutants (Rohner et al., 2009), suggesting that the severity of this phenotype may be influenced by the genetic background. Alternatively, it is possible that *fgfr1a(t3R05H*) is a weak anti-morph mutation, as it is known that over-expression of an Fgfr lacking a functional intercellular kinase domain can be dominant-negative (Amaya et al., 1991; Griffin et al., 1995).

### Genetic redundancy and transcriptional compensation in Fgf receptor mutants

Recently, it has been shown that indel alleles generated by genome editing technologies can result in phenotypes that are weaker than either point mutant alleles or morpholino oligo knockdown. This is likely due to a mechanism known as genetic compensation, where the transcription of a gene(s) related to the mutated gene is upregulated in mutants and functionally compensates, either partially or completely, for the mutated gene (Rossi et al., 2015). The mechanism by which transcriptional compensation occurs is not currently known. Given that the Fgf receptors share extensive sequence similarity, it was possible that transcriptional compensation could account for the lack of phenotype in our Fgfr mutants. To test this, we used reverse transcription quantitative real-time PCR (RT-qPCR) to examine the expression of all *fgfr* genes in our *fgfr* mutants. We compared *fgfr* gene expression between individual wild-type, single mutant and select double and triple mutant 24 hours post fertilization (hpf) embryos (Fig. 2). Although expression changes are modest, trends emerge from our data. First, with one exception, the mRNA of the mutated gene is detected at significantly lower levels compared to wild-type, suggesting that the mutant mRNAs are subject to nonsense-mediated decay. The exception to this is that, in comparison to wild-type controls, *fgfr2* mRNA is significantly decreased in *fgfr2*^-/-^ single (Fig. 2C) and *fgfr1a*^-/-^*;fgfr2*^-/-^ double mutants (Fig. 2G), but not in *fgfr1a*^-/-^*;fgfr1b*^-/-^*;fgfr2*^-/-^ triple mutant embryos (Fig. 2H; Fig. 2I shows confirmation that these triple mutant embryos are indeed mutant for *fgfr2*). The reason for this is not known.

**Fig. 2.**
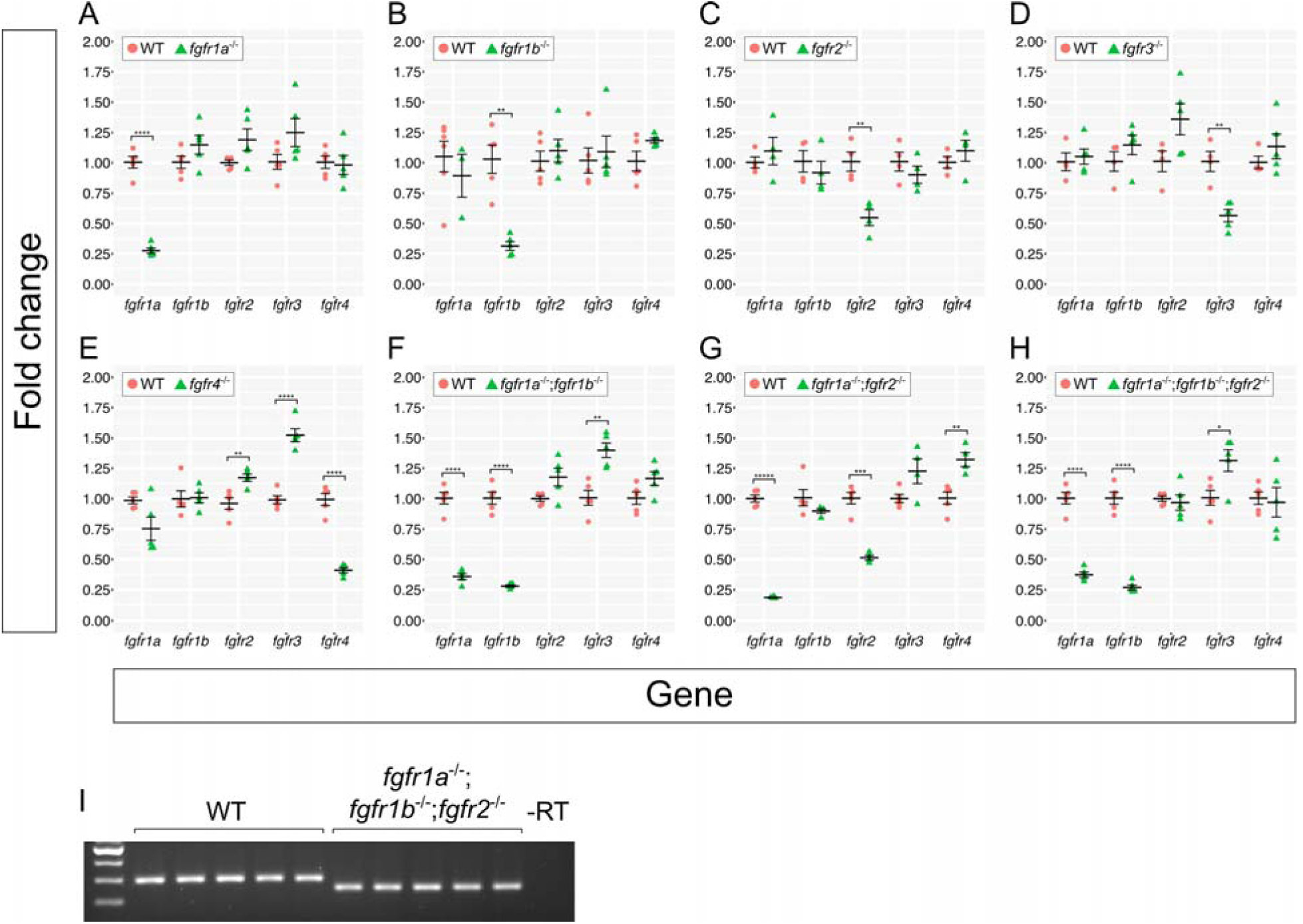
Fgf receptor mRNAs are overexpressed in some Fgf receptor mutants compared to wild-type embryos. (A-H) Fold changes calculated from RT-qPCR experiments comparing Fgf receptor mRNA levels between wild-type (WT, pink circles) and various Fgf receptor mutants (green triangles). Each point represents the mean fold change of an individual embryo, relative to WT. Error bars represent ± the standard error of the mean (center bar). Number of asterisks represents p-values calculated using a Student’s T-test: no asterisk between WT and mutant denotes p > 0.05; *p < 0.05; **p < 0.01; ***p < 0.001; ****p < 0.0001; *****p < 0.00001. (I) Because the *fgfr1a*^-/-^*;fgfr1b*^-/-^*;fgfr2*^-/-^ triple mutants represented in (H) displayed near-WT levels of *fgfr2* mRNA, *fgfr2* genotypes were confirmed with standard RT-PCR. Note the decreased band size in all *fgfr1a;fgfr1b;fgfr2*^-/-^ mutants.

Second, with the exception of *fgfr4* mutants, which have a significant increase in the expression of wild-type *fgfr2* and *fgfr3* RNA (Fig. 2E), none of the other single mutants had significant increases in the expression of any of the wild-type *fgfr* RNAs (Fig. 2A-D). By contrast, all double and triple mutant combinations we examined had significantly higher levels of wild-type *fgfr3* mRNA (*fgfr1a*^-/-^;*fgfr1b*^-/-^ and *fgfr1a*^-/-^;*fgfr1b*^-/-^;*fgfr2*^-/-^) or *fgfr4* (*fgfr1a*^-/-^;*fgfr2*^-/-^) mRNA in comparison to wild-type controls (Fig. 2F-H). Whether these increases in wild-type *fgfr* RNA expression in the mutants result in less severe terminal phenotypes remains to be determined. Regardless, these data argue that the lack of phenotypes in most of the <u>single</u> *fgfr* mutants is not likely due to genetic compensation, but instead is due to genetic redundancy.

### *fgfr1a* and *fgfr1b* are required for posterior mesoderm development

In vertebrates, the posterior body derives from the tailbud, a population of multipotent cells that forms at the caudal end of the embryo at the end of gastrulation. These cells undergo proliferation and differentiation resulting in the elongation of the body axis in the posterior direction. Lineage analyses have shown that the tail bud gives rise to the hindgut, posterior neural tube, notochord, and somites. Fgf signaling is known to play a role in posterior mesoderm development; both *Fgfr1* and *Fgf8* mouse mutants lack posterior mesoderm due to failures in mesodermal specification and morphogenesis at the primitive streak (Ciruna and Rossant, 2001; Deng et al., 1994; Sun et al., 1999; Yamaguchi et al., 1994). In zebrafish, animals expressing a dominant-negative Fgfr form no posterior body (Griffin et al., 1995), whereas animals deficient for both *fgf8a* and *fgf24* have a somewhat less severe reduction of posterior mesoderm (Draper et al., 2003). In this latter study, it was shown that *fgf8a* and *fgf24* are together required to maintain, but not initiate, the expression of *ta* (*ntl*/*brachyury homolog a*) and *tbx16* (*spt*), T-box transcription factor genes known to be required for mesodermal specification (Amacher et al., 2002; Conlon et al., 1996; Halpern et al., 1993; Kimmel et al., 1989; Warga et al., 2013; Zhang et al., 1998). These expression defects are visible around 80% epiboly (Draper et al., 2003), a time at which only *fgfr1a* and *fgfr1b* are expressed highly at the margin of the gastrulating embryo where mesodermal precursors reside (Rohner et al., 2009). By contrast, during gastrulation, *fgfr2* and *fgfr3* have minimal expression in mesodermal precursors and *fgfr4* expression appears to be restricted to cells that reside closer to the animal pole (Ota et al., 2010). These expression data therefore identify *fgfr1a* and *fgfr1b* as the likely candidate receptors involved in posterior mesoderm development.

Previously, Rohner et al. showed a variable posterior defect in animals homozygous for *fgfr1a(t3R05H*) that were also injected with morpholinos targeting *fgfr1b* (Rohner et al., 2009). We therefore asked whether our *fgfr1a*;*fgfr1b* double mutant animals had defects in posterior mesoderm development. Although these double mutants die around 5 days post fertilization (dpf), we found that in comparison to wild-type animals (Fig. 3A), *fgfr1a;fgfr1b* double mutant animals have shorter and slightly kinked tails and an accumulation of blood on the ventral side posterior to the yolk extension (Fig. 3B). Because both *fgfr1a* and *fgfr1b* mRNAs are maternally provided (Fig. 1C), it was possible that the mild defects observed were due to maternal gene product. To test this, we produced various combinations of maternal and/or zygotic loss of *fgfr1a* and *fgfr1b*. We found that Z*fgfr1*a;MZ*fgfr1b* mutants were largely indistinguishable from Z*fgfr1a*;Z*fgfr1b* mutants (Fig. 3B,C), arguing that maternally provided *fgfr1b* was not responsible for the mild phenotype. By contrast, we found that MZ*fgfr1a*;Z*fgfr1b* embryos had significantly shorter tails than Z*fgfr1a*;Z*fgfr1b* mutant embryos (Fig. 3B,D). These data argue that normal posterior mesoderm development requires zygotically-expressed *fgfr1a* and *fgfr1b*, but also maternally-expressed *fgfr1a*. Because *fgfr2* appears to be redundant with *fgfr1a* and *fgfr1b* in the other developmental contexts reported here (Figs 4-7), we also asked whether the additional removal of functional *fgfr2* from Z*fgfr1a*;Z*fgfr1b* or MZ*fgfr1a*;Z*fgfr1b* double mutant embryos enhanced the respective phenotypes. However, these triple mutants underwent similar posterior mesoderm development to their double mutant counterparts, suggesting that *fgfr2* is not required for this process (Fig. S1B,C).

**Fig. 3.**
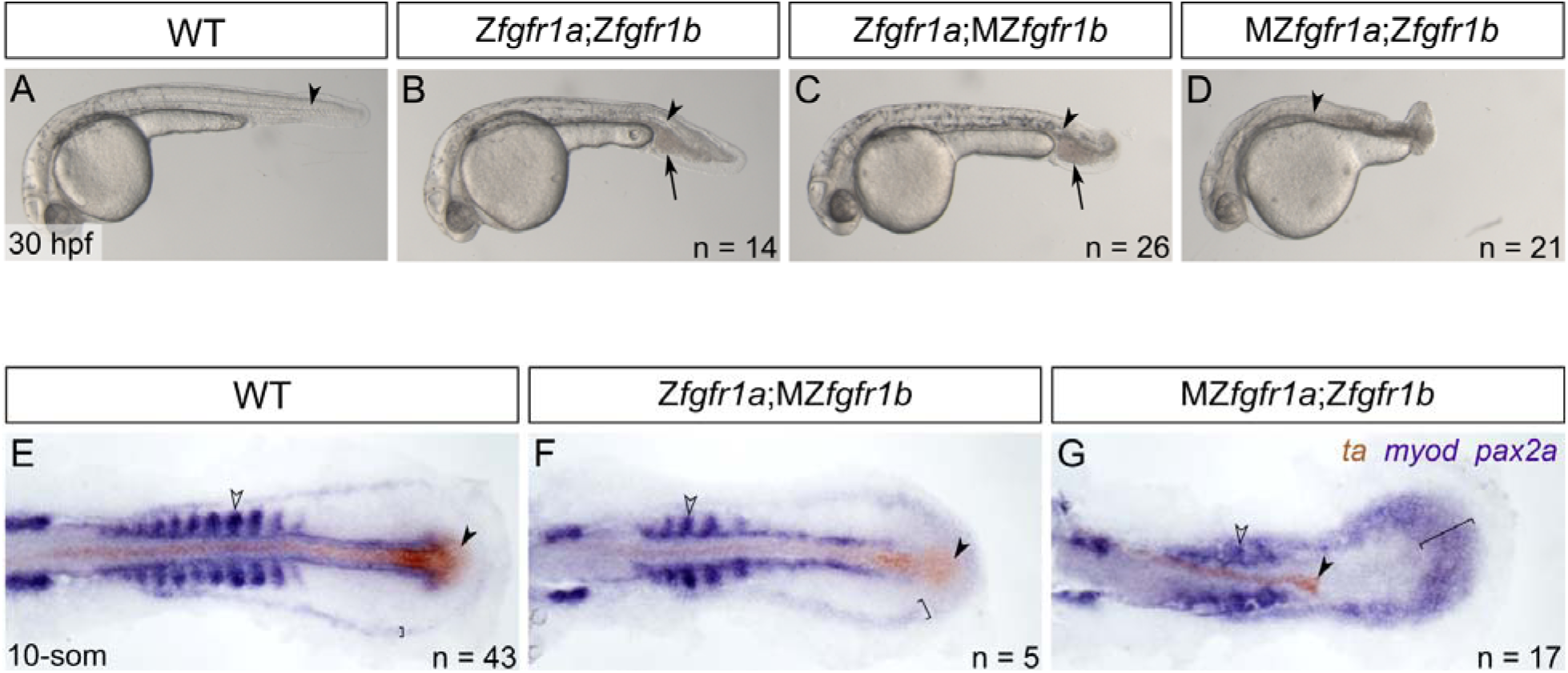
*fgfr1a* and *fgfr1b* function redundantly to regulate posterior mesoderm development. (A-D) Lateral view of 30 hpf wild-type (WT; A), Z*fgfr1a*;Z*fgfr1b* (B), Z*fgfr1a*;MZ*fgfr1b* (C), and MZ*fgfr1a*;Z*fgfr1b* (D) mutant embryos. Arrowheads denote the notochord; arrows mark pooled blood cells in B and C. Anterior is to the left, dorsal is up. (E-G) Mesodermal derivative marker analysis of Z*fgfr1a*;MZ*fgfr1b* and MZ*fgfr1a*;Z*fgfr1b* double mutant embryos at the 10-somite stage. The notochord (labeled with *ta*, brown; filled arrowhead) extends down the length of the posterior body in wild-type (E) and Z*fgfr1a*;MZ*fgfr1b* double mutant embryos (F) but is truncated in MZ*fgfr1a*;Z*fgfr1b* double mutant embryos (G). Defined somites (labeled with *myod*, purple; open arrowhead) are present in wild-type embryos (E) and Z*fgfr1a*;MZ*fgfr1b* embryos (F), however the latter have distinctly fewer somites (5 compared to 10). Although MZ*fgfr1a*;Z*fgfr1b* mutants retain some *myod*-positive cells, there are no definitive somites (G). Pronephric precursors (labeled with *pax2a*, purple; brackets) are restricted to a defined band around the posterior body of wild-type embryos (E), a region that is expanded in both Z*fgfr1a*;MZ*fgfr1b* (F) and MZ*fgfr1a*;Z*fgfr1b* double mutant embryos (G).

**Fig. 4.**
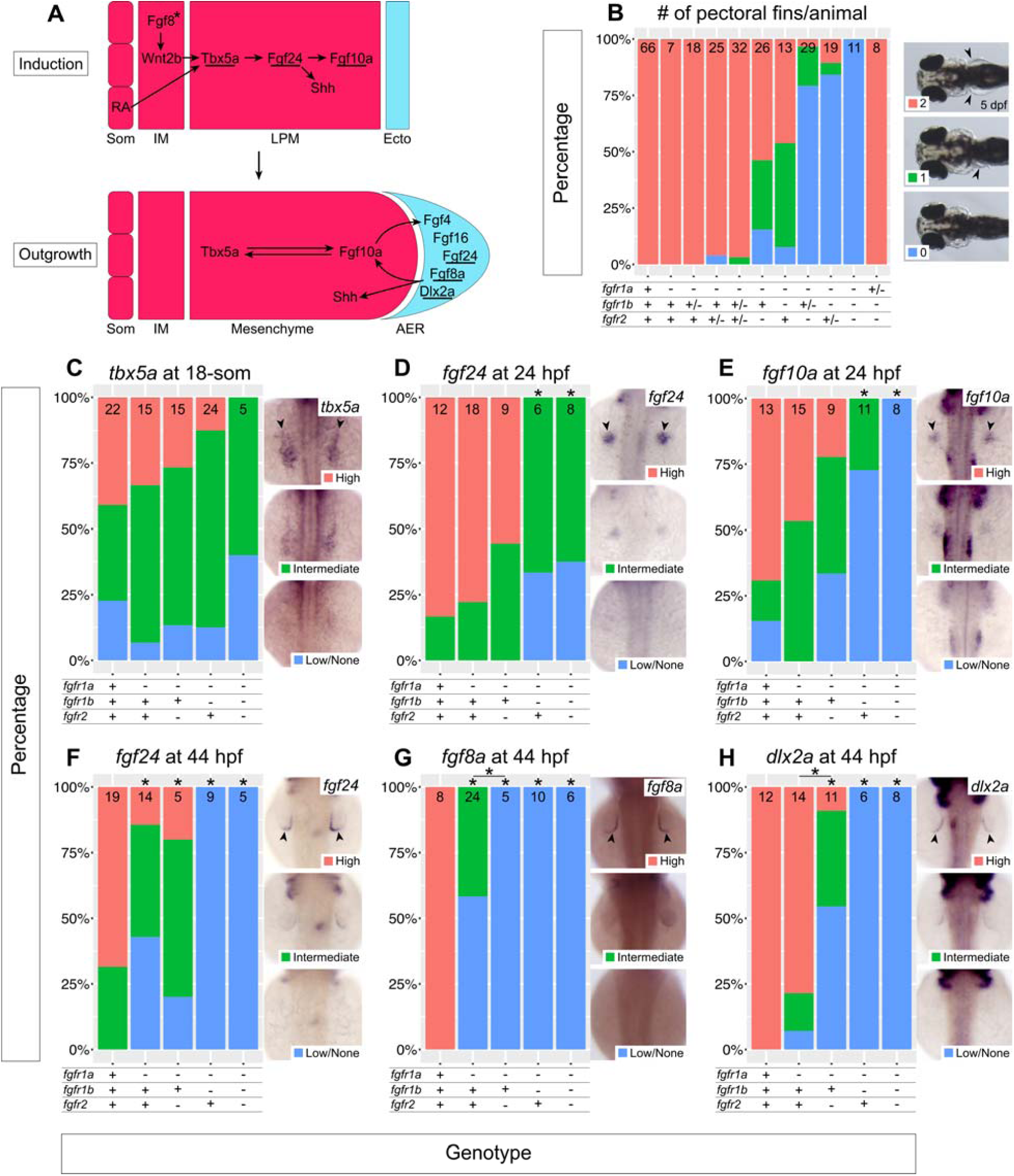
*fgfr1a, fgfr1b*, and *fgfr2* function redundantly to regulate pectoral fin development. (A) Model of vertebrate forelimb bud development during forelimb bud Induction (top), Outgrowth (bottom). Underlines denote genes assayed in (C-H). Arrows denote an epistatic (but not direct) link between molecules. Asterisk signifies that Fgf8 has not yet been shown to play this role in zebrafish, but is hypothesized from work in chick and mouse. (B) Stacked column chart depicting the average number of pectoral fins per animal at 5 dpf, according to genotype. Sample size for each genotype is listed at the top of each bar. Representative images of larvae with 2, 1, or 0 pectoral fins to the right: dorsal views, anterior to the left, with arrowheads denoting pectoral fins where present. (C-H) Limb bud marker analysis of *fgfr* double and triple mutant embryos at the 18-somite stage (*tbx5a*, C), 24 hpf (*fgf24*, D; *fgf10a*, E), and 44 hpf (*fgf24*, F; *fgf8a*, G; *dlx2a*, H). Whole mount *in situ* hybridization was performed, embryos were scored for expression, and genotypes were determined post-hoc. In each panel, the percentage of embryos expressing particular levels of each marker gene is represented in a stacked column chart on the left, and representative images of those expression levels are shown for each marker to the right (dorsal views, anterior up; developing limb buds are seen as two spots on either side of the embryo, denoted by arrowheads).

To further examine the posterior defects, we compared gene expression between wild-type, Z*fgfr1a*;MZ*fgfr1b*, and MZ*fgfr1a*;Z*fgfr1b* 10-somite stage embryos by RNA *in situ* hybridization to assess the relative amounts of mesodermal derivatives produced. We co-stained embryos for the axial mesoderm marker *ta* (formerly *ntl*), which labels notochord cells (Schulte-Merker et al., 1992), and the paraxial mesodermal marker *myod*, which labels somitic mesoderm (Weinberg et al., 1996). We found that while wild-type and Z*fgfr1a*;MZ*fgfr1b* embryos have a notochord that extends down the entire length of the trunk and tail, MZ*fgfr1a;*Z*fgfr1b* mutant embryos formed notochord only in the trunk region (filled arrowheads, Fig. 3E-G). Similarly, we found that at 14 hpf, when wild-type embryos had produced 10 somites, Z*fgfr1a*;MZ*fgfr1b* embryos have formed only five somites (Fig. 3E,F). Finally, in MZ*fgfr1a*; Z*fgfr1b* mutant embryos, although somatic mesoderm appears to have formed based on *myod* expression, proper somite morphogenesis appears to have failed (Fig. 3G; open arrowhead). These findings are similar to that of *fgf8a* mutants injected with *fgf24* morpholino (MO)s, suggesting that these ligands signal, at least in part, via Fgfr1a and Fgfr1b during posterior mesoderm development. However MZ*fgfr1a*;Z*fgfr1b* mutants form more posterior mesoderm than *fgf8a* mutant; *fgf24*MO animals (Draper et al., 2003), and significantly more than animals expressing a dominant-negative Fgfr (Griffin et al., 1995). We therefore hypothesize that residual activity of one or more of the remaining three Fgf receptors (Fgfr2, Fgfr3, Fgfr4) must be sufficient to promote partial production of posterior tissue.

*pax2a* is a marker of pronephric precursors, and in wild-type embryos is restricted to a discrete domain of intermediate mesoderm in the 10-somite stage embryo (bracket, Fig. 3E; (Draper et al., 2003; Krauss et al., 1991)). By contrast, Z*fgfr1a*;MZ*fgfr1b* and, to a greater extent, MZ*fgfr1a*;Z*fgfr1b* mutants have an expanded region of *pax2a*-expressing cells, suggesting that these mutants produce an increased number of pronephric precursors relative to wild-type embryos (brackets, Fig. 3F,G). Additionally, Z*fgfr1a*;Z*fgfr1b* and Z*fgfr1a*;MZ*fgfr1b* mutants have an accumulation of blood cells just posterior to the yolk extension, similar to dorsalized mutants, such as *chordin* (arrows, Fig. 3B,C; (Wagner and Mullins, 2002)). Given that the pronephros and blood are both derived from intermediate mesoderm, and somites and notochord arise from more dorsal paraxial and axial mesoderm, respectively, these data are consistent with previous analyses of *fgf8a* mutants, which concluded that Fgf signaling is important for promoting dorsal mesodermal fates (Furthauer et al., 1997; Furthauer et al., 2004).

### *fgfr1a, fgfr1b* and *fgfr2* are required for pectoral fin development

Pectoral fins are the equivalent of mammalian forelimbs, and, with few exceptions, their development is regulated by orthologs of genes that regulate mammalian forelimb development. Studies using mouse, chick, and zebrafish have identified the important and conserved signaling ligands that participate in this process (Fig. 4a). For example, in both forelimb and pectoral fin development, the limb field within the lateral plate mesoderm (LPM) is specified by signals produced by the paraxial (retinoic acid) and intermediate mesoderm (FGF8 and WNT2B), which initiates expression of the T-box transcription factor gene, *TBX5/tbx5a*, in the LPM (Cohn et al., 1995; Crossley et al., 1996b; Gibert et al., 2006; Kawakami et al., 2001; Ng et al., 2002). Subsequently, TBX5/Tbx5a activates the expression of an Fgf ligand, *Fgf10*/*fgf10a* within the LPM (in zebrafish, this step is mediated by another Fgf ligand, Fgf24) (Fischer et al., 2003; Min et al., 1998; Ng et al., 2002). FGF10/Fgf10a in turn acts upon the overlying ectoderm to initiate formation of the apical ectodermal ridge (AER), a structure that reciprocally signals back to the mesoderm via additional Fgf ligands (e.g. FGF2, FGF4, FGF8 in mouse and chick; Fgf4, Fgf8a, Fgf24, and Fgf16 in zebrafish; (Crossley et al., 1996b; Fallon et al., 1994; Fischer et al., 2003; Kawakami et al., 2001; Kengaku et al., 1998; Laufer et al., 1994; Min et al., 1998; Niswander and Martin, 1992; Nomura et al., 2006; Sun et al., 2002)) to maintain the expression of limb development genes within the fin bud mesenchyme. This Fgf-dependent feedback loop is maintained for the duration of limb development and is required for limb outgrowth and patterning along the proximodistal axis (reviewed in (Xu et al., 1999b)). Thus, limb development requires a complex signaling network, of which Fgf signaling is a key component, to coordinate its development.

In mouse, null mutations of *Fgfr1* and *Fgfr2* result in embryonic lethality before the end of gastrulation, precluding their use for determining their role in limb development. However, the use of hypomorphic alleles have led to the conclusion that FGFR1 is involved in limb patterning, while FGFR2 has a more prominent role in limb bud induction and outgrowth (De Moerlooze et al., 2000; Xu et al., 1999a; Xu et al., 1998), (Xu et al., 1999b). Given these findings and the established roles for Fgf ligands throughout limb development, it was surprising that all five single *fgfr* mutants have normal pectoral fin development (arrowheads, Fig. 4B, and data not shown). However, *a*; *b a* at 5 dpf, we found that 54% of *fgfr1a* ^-/-^ *;fgfr1* ^-/-^ double mutants and 46% of *fgfr1a* ^-/-^;*fgfr2* ^-/-^ double mutants lack at least one pectoral fin (n = 13 and n = 26, respectively), establishing a role for all three of these receptors in pectoral fin development. Removing the function of an additional *fgfr* allele in these double homozygous mutants increases the penetrance of the pectoral fin phenotype: 90% of *fgfr1a* ^-/-^ *fgfr1b* ^-/-^ *fgfr2* ^+/-^ and 97% of *fgfr1a* ^-/-^ *fgfr1b* ^+/-^ *fgfr2* ^-/-^ lack at least one pectoral fin (n = 19 and n = 29, respectively). Finally, 100% of *fgfr1a* ^-/-^ *fgfr1b* ^-/-^ *fgfr2* ^-/-^ triple mutants fail to form any pectoral fins (n = 11; Fig. 4B), indicating that these three receptors act redundantly to promote zebrafish pectoral fin development. Interestingly, the presence of a single wild-type copy of *fgfr1a* is sufficient to rescue this phenotype completely (Fig. 4B), suggesting that *fgfr1a* is particularly important for pectoral fin development.

In zebrafish, pectoral fin bud initiation occurs around the 18-somite stage (18 hpf), as evident by the expression of *tbx5a*, one of two orthologs of mouse and chick *Tbx5* (Ahn et al., 2002; Garrity et al., 2002; Ng et al., 2002). Shortly thereafter, Tbx5a promotes the transcription of *fgf24* within the fin bud mesoderm, which is then required for mesodermal transcription of *fgf10a* and *sonic hedgehog* (*shh*; involved in anterior-posterior patterning of limbs) (Fischer et al., 2003; Krauss et al., 1993; Krauss et al., 1991; Neumann et al., 1999; Ng et al., 2002). Fgf10a maintains *tbx5a* expression in the fin bud mesoderm and is likely responsible for signaling to the overlying ectoderm to induce AER formation, which includes the induction of *fgf8a, fgf4, fgf24*, and *fgf16* expression in the ectoderm by ∼30-36 hpf (Fischer et al., 2003; Grandel et al., 2000; Ng et al., 2002; Nomura et al., 2006; Reifers et al., 1998). In chicken and mice this induction appears to be mediated by WNT signaling (Kawakami et al., 2001; Kengaku et al., 1998), however this has yet to be established in zebrafish. Similar to chick and mice, it is likely that the AER Fgfs signal back to the fin bud mesenchyme to stabilize *fgf10a* expression, thus establishing a positive regulatory feedback loop required for fin bud outgrowth (Fig. 4A) (Camarata et al., 2010).

Because Fgf signaling is known to mediate many tissue interactions during limb development, we sought to characterize at what level the various mutant combinations affect fin development by assessing the expression of marker genes using RNA *in situ* hybridization at different stages of fin bud development. Following *in situ* hybridization, embryos were scored for marker gene expression first and then genotyped. We initially asked if any of the mutant combinations affected fin bud initiation by assaying the expression of *tbx5a*, the earliest marker of pectoral fin bud induction (Ahn et al., 2002). In wild-type embryos, *tbx5a* expression in the LPM can be detected in most, but not all, 18-somite stage embryos, a stage that precedes feedback regulation from the LPM expressed Fgfs (Fig. 4A,C). Expression was similarly detected in all other genotypes examined, including *fgfr1a*^-/-^;*fgfr1b*^-/-^;*fgfr2*^-/-^ triple mutants, though the triple mutants had, on average, less intense staining (Fig. 4C). These results suggest that the first steps of limb bud induction are only mildly affected by loss of Fgfr1a, Fgfr1b, and Fgfr2 function (arrowheads, Fig. 4C).

In zebrafish, Tbx5a induces the expression of *fgf24* in the LPM, and Fgf24 signals within the LPM to stimulate *fgf10a* expression, which in turn is thought to form a feedback loop to maintain *tbx5a* expression (Fig. 4A)(Fischer et al., 2003; Ng et al., 2002). While the expression of *fgf24* and *fgf10a* is easy to detect by *in situ* hybridization in most 24 hpf wild-type fin buds, their expression is reduced or not detected in 24 hpf *fgfr1a;fgfr1b* and *fgfr1a;fgfr2* double mutant and *fgfr1a;fgfr1b;fgfr2* triple mutant fin buds (Fig. 4D,E). These results suggest that, although the fin bud is induced in these mutants, Fgfr1a, Fgfr1b, and Fgfr2 are redundantly required to maintain gene expression within the limb bud mesenchyme.

During the outgrowth phase of limb development (24 hpf - ∼48 hpf) (Grandel and Schulte-Merker, 1998), Fgfs from the fin bud mesenchyme signal the ectoderm to form the AER. We therefore asked if AER-specific gene expression was reduced in *fgfr1a;fgfr1b* and *fgfr1a;fgfr2* double mutants and *fgfr1a;fgfr1b;fgfr2* triple mutants. Indeed, whereas *fgf24, fgf8a*, and *dlx2a* are all expressed in the AER of wild-type animals at 44 hpf, the number of embryos with reduced or no detectable expression by *in situ* hybridization is greatly increased in the various mutant combinations (Fig. 4F-H). Interestingly, in *fgfr1a* single mutant embryos, which have normal pectoral fin development, we also observed a reduction in AER-expressed *fgf24* and *fgf8a* at 44 hpf (Fig. 4F). Together these results suggest that Fgfr1a, Fgfr1b, and Fgfr2 function redundantly to maintain proper gene expression in the limb bud mesenchyme and subsequently in the AER, but that *fgfr1a* may be particularly important at the mesenchyme/AER interface, and may explain why the presence of a single wild-type copy of *fgfr1a* is sufficient to rescue the *fgfr1a;fgfr1b;fgfr2* triple mutant pectoral fin phenotype (Fig. 4B).

### *fgfr1a, fgfr1b* and *fgfr2* are required for brain development

The vertebrate brain develops from the relatively simple neural plate. One of the earliest patterning events of the neural plate is its subdivision into rostral and caudal domains that can be identified by expression of the transcription factors *Otx2* and *Gbx2*, respectively (Broccoli et al., 1999; Millet et al., 1999). A signaling center called the midbrain-hindbrain organizer forms at the boundary of these domains, which acts to pattern the surrounding neural tissues (Marin and Puelles, 1994; Martinez et al., 1995). Fgf signaling is the most prominent signaling pathway in the MHB, and while many Fgf ligands are known to be expressed in the MHB organizer (Fgf8, Fgf17, Fgf18, Fgf4), Fgf8 appears to have the most critical role: in chick, FGF8-soaked beads ectopically induce midbrain development (Crossley et al., 1996a), whereas mutations in mouse *Fgf8*, or its ortholog *fgf8a* in zebrafish, result in loss of the MHB and cerebellum (Brand et al., 1996; Reifers et al., 1998) (Chi et al., 2003; Meyers et al., 1998).

In mice, tissue-specific knock-out of *Fgfr1* in MHB cells, or a *Fgfr1* hypomorphic mutation, leads to the loss of certain MHB structures. However, this phenotype is less severe than that of *Fgf8* mutants, suggesting that other receptors are involved in FGF8 signal transduction during MHB development (Chi et al., 2003; Trokovic et al., 2003). In zebrafish, morpholino knock-down of *fgfr1a* has been reported to phenocopy *fgf8a* mutants (Scholpp et al., 2004). By contrast to the *fgfr1a* morphants, *fgfr1a(uc61)* and *fgfr1a(t3R05H*) mutants have normal MHB development, strongly arguing that in zebrafish, Fgf receptors in addition to Fgfr1a are able to mediate Fgf8a signaling during MHB development. We therefore produced double and triple mutant combinations to test if additional receptors are involved in MHB development.

Zebrafish *fgf8a* mutants first display gene expression abnormalities in the hindbrain region during early somitogenesis, with increasing severity by late somitogenesis (18-somite stage) (Reifers et al., 1998). At the 18-somite and 24 hpf stages, when the MHB signaling center is active (Ota et al., 2010), all five Fgf receptor genes can be detected in the MHB region (Ota et al., 2010; Rohner et al., 2009; Scholpp et al., 2004; Thisse, 2001, 2008; Thisse, 2005; Tonou-Fujimori et al., 2002). However, considering the apparent role for FGFR1 in maintenance of midbrain and hindbrain tissue in the mouse, we first tested if *fgfr1a*;*fgfr1b* double mutants would lead to a brain defect similar to that of the *fgf8a* mutation. To our surprise, the MHBs of these animals are morphologically indistinguishable from wild-type (Figs. 5A, B). In contrast, the triple mutant *fgfr1a*;*fgfr1b*;*fgfr2* appears to phenocopy the acerebellar phenotype of *fgf8*a mutants. (Fig. 5C).

**Fig. 5.**
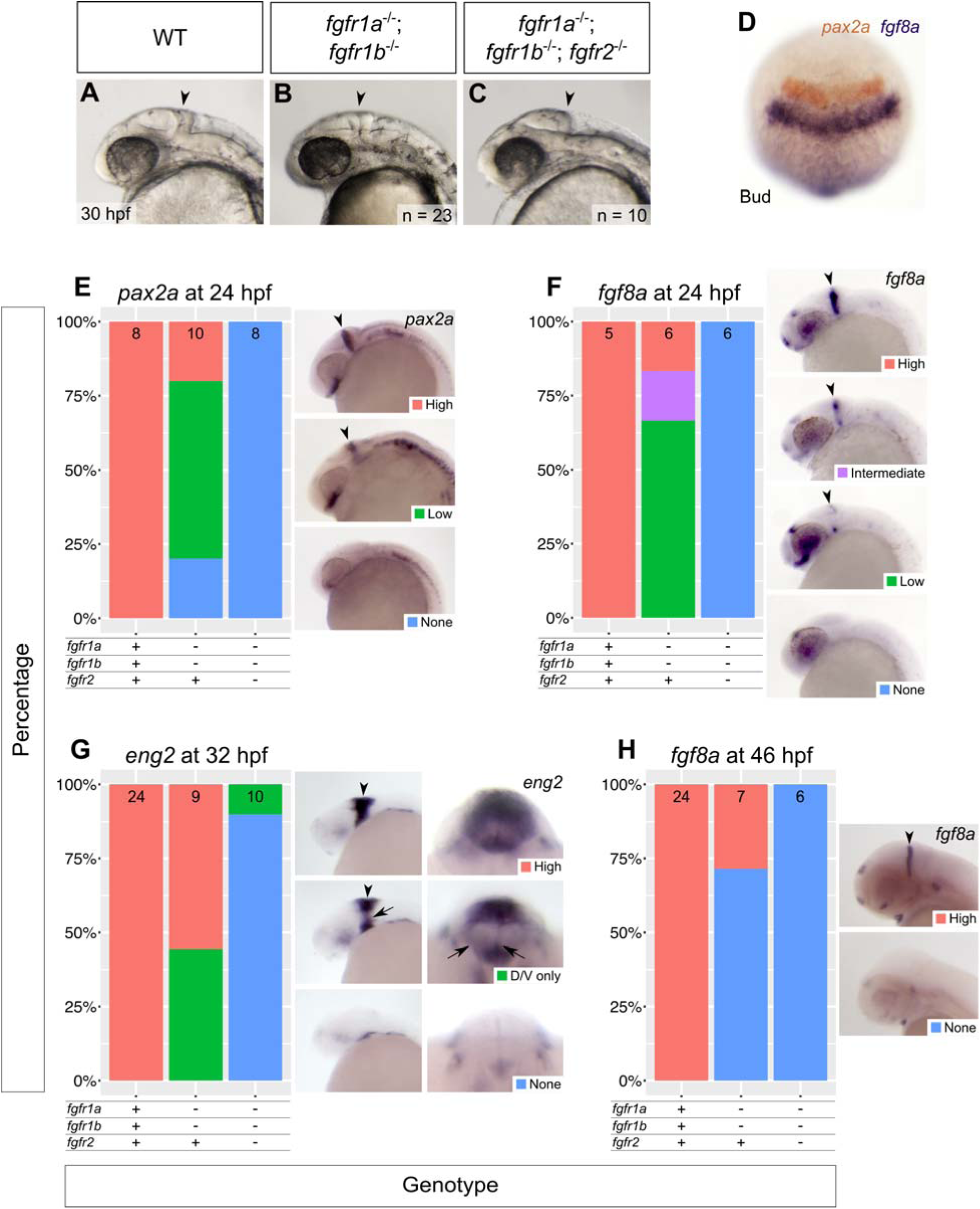
*fgfr1a, fgfr1b*, and *fgfr2* function redundantly to regulate MHB development. (A-C) Lateral view of 30 hpf wild-type (WT, A), *fgfr1a*^-/-^;*fgfr1b*^-/-^ (B; n = 23), *fgfr1a*^-/-^;*fgfr1b*^-/-^;*fgfr2*^-/-^ (C; n = 10) mutant embryos. Arrowheads denote region where the MHB should form. Rostral is to the left, dorsal is up. (D-H) MHB marker analysis of *fgfr* double and triple mutant embryos at the bud stage (*pax2a* in brown/*fgf8a* in purple, D), 24 hpf (*pax2a*, E; *fgf8a*, F), 32 hpf (*eng2*, G), and 46 hpf (*fgf8a*, H). Whole mount *in situ* hybridization was performed, embryos were scored for expression, and genotypes were determined post-hoc. All embryos had indistinguishable *pax2a*/*fgf8a* expression at the bud stage (D; n = 28, 6, 4, for WT, *fgfr1a*^-/-^;*fgfr1b*^-/-^, and *fgfr1a*^-/-^;*fgfr1b*^-/-^;*fgfr2*^-/-^, respectively). In (E-H), the percentage of embryos expressing particular levels of each marker gene is represented in a stacked column chart on the left (sample size for each genotype is listed at the top of each bar), and representative images of those expression levels are shown for each marker to the right (lateral views, rostral to the left and dorsal up; developing MHBs are denoted by arrowheads). Rightmost images in (G) are frontal views (dorsal up) showing low *eng2* staining in the left and right regions of the cerebellum (arrows).

To further assess the brain development of these animals, we used RNA *in situ* hybridization to assay the expression of genes known to play a role in MHB development. At the bud stage (10 hpf) of development, *pax2a* and *fgf8a* label the prospective MHB (Krauss et al., 1991; Mikkola et al., 1992; Reifers et al., 1998). In all genotypes examined, including *fgfr1a;fgfr1b;fgfr2* triple mutants, we found that both genes are expressed in their normal locations, suggesting that the specification of MHB cells is unaffected (Fig. 5D). In contrast, at 24 hpf, when both *pax2a* and *fgf8a* are still expressed in the MHB of wild-type embryos, their expression is not detected in the MHB of *fgfr1a;fgfr1b;fgfr2* triple mutants (Fig. 5E,F). Consistent with this, the cerebellar marker *en2a* is absent from most triple mutants by 32 hpf. (Fig. 5G,H) (Millen et al., 1994).

Surprisingly, although *fgfr1a*;*fgfr1b* double mutants have apparent wild-type brain morphology at 24 hpf (Fig. 5B), we found that marker gene expression was reduced in many double mutant embryos when compared to wild-type animals at 24 hpf, and by 46 hpf, *fgf8a* expression is not detectable in ∼70% of double mutant embryos (Fig. 5E-H). These data suggest that a partial reduction in Fgf signaling through Fgfr1a is sufficient to affect gene expression in the MHB, but not overt MHB morphology. Though it is possible that brain patterning, and therefore brain function, is compromised in these animals, we have not been able to test this as they do not survive past 5 dpf.

### *fgfr1a, fgfr1b* and *fgfr2* are required for craniofacial skeletal development

Skeletal structures of the head are largely derived from cranial neural crest cells. During development, neural crest cells (NCCs) migrate ventrolaterally from the dorsal neuroectoderm of the hindbrain into endodermal pockets called pharyngeal pouches. There, intrinsic cues and inductive signaling from the surrounding tissue instruct NCC development into cartilage (reviewed in (Kimmel et al., 2001; Mork and Crump, 2015). In zebrafish, seven arches form between the endodermally-derived pouches, and most arch derivatives compose the viscerocranium: the first arch gives rise to Meckel’s cartilage and the palatoquadrate; the second gives rise to the ceratohyal and hyosemplectic; and arches 3-7 give rise to the ceratobranchials (Crump et al., 2006; Crump et al., 2004b; Kimmel et al., 2001; Schilling and Kimmel, 1994). However, lineage tracing has revealed that NCCs also give rise to the neurocranium; NCCs from the first two arches contribute to discrete portions of the postchordal neurocranium, whereas the prechordal neurocranium arises from more anteriorly-derived NCCs (Eberhart et al., 2006; McCarthy et al., 2016; Swartz et al., 2011; Wada et al., 2005).

Many Fgfs and their receptors are expressed throughout the head during the time of cranial cell specification and differentiation (Larbuisson et al., 2013; Ota et al., 2010), and several of the processes underlying cranial morphogenesis are known to be driven by Fgf signaling. In mice, *Fgf8* is required for proper development of several pharyngeal arch-derived craniofacial structures, and FGFR1 and FGFR2 play an important role in mammalian palatogenesis (Abu-Issa et al., 2002; Rice et al., 2004; Yu et al., 2015). In zebrafish, *fgf8a*;*fgf3* double mutants do not form the posterior viscerocranium, have severely deformed anterior viscerocranium, and appear to not form the mesodermally-derived cartilages of the postchordal neurocranium (Crump et al., 2004a; McCarthy et al., 2016). Furthermore, morpholino analysis suggests that the Fgf receptors Fgfr1a and Fgfr2 can each regulate late cartilage formation in the viscerocranium, although the effects are attributed to later defects compared to that caused by loss of *fgf8a* and *fgf3* (Larbuisson et al., 2013). However, we found that cranial development in *fgfr1a* and *fgfr2* single mutants, as well as *fgfr1a; fgfr2* double mutants were indistinguishable from wild-type (data not shown). By contrast, we found that *fgfr1a;fgfr1b* double mutants have reduced viscerocrania, including a loss of most of the ceratobranchials and hyosymplectic, and also have misshapen palatoquadrate and Meckel’s cartilage (Fig. 6B-B’’). Although *fgfr1a;fgfr1b* double mutant animals that are also heterozygous for a null allele of *fgfr2* (i.e. *fgfr1a*^-/-^;*fgfr1b*^-/-^;*fgfr2*^+/-^ are indistinguishable from *fgfr1*^-/-^;*fgfr1b*^-/-^*fgfr2*^+/+^ animals with respect to viscerocrania development (Fig. 6B-B”), both *fgfr1a*^-/-^;*fgfr1b*^-/-^;*fgfr2*^+/-^ and *fgfr1a*^-/-^;*fgfr1b*^-/-^;*fgfr2*^-/-^ mutants lack portions of the postchordal neurocranium that are thought to be derived from the first and second pharyngeal arches (Fig. 7C,C’). Finally, of all genotypes analyzed, *fgfr1a;fgfr1b;fgfr2* triple mutants have the most severe cranial defects: in addition to an abnormal postchordal neurocranium (Fig. 7C,C’), the viscerocranial ceratobranchials, ceratohyal, and most of the hyosymplectic fail to form, the palatoquadrate is misshapen, and Meckel’s cartilage is displaced downward (Fig. 6C-C’’). Together, these data suggest that Fgfr1a, Fgfr1b, and Fgfr2 function redundantly to promote cranial cartilage formation.

**Fig. 6.**
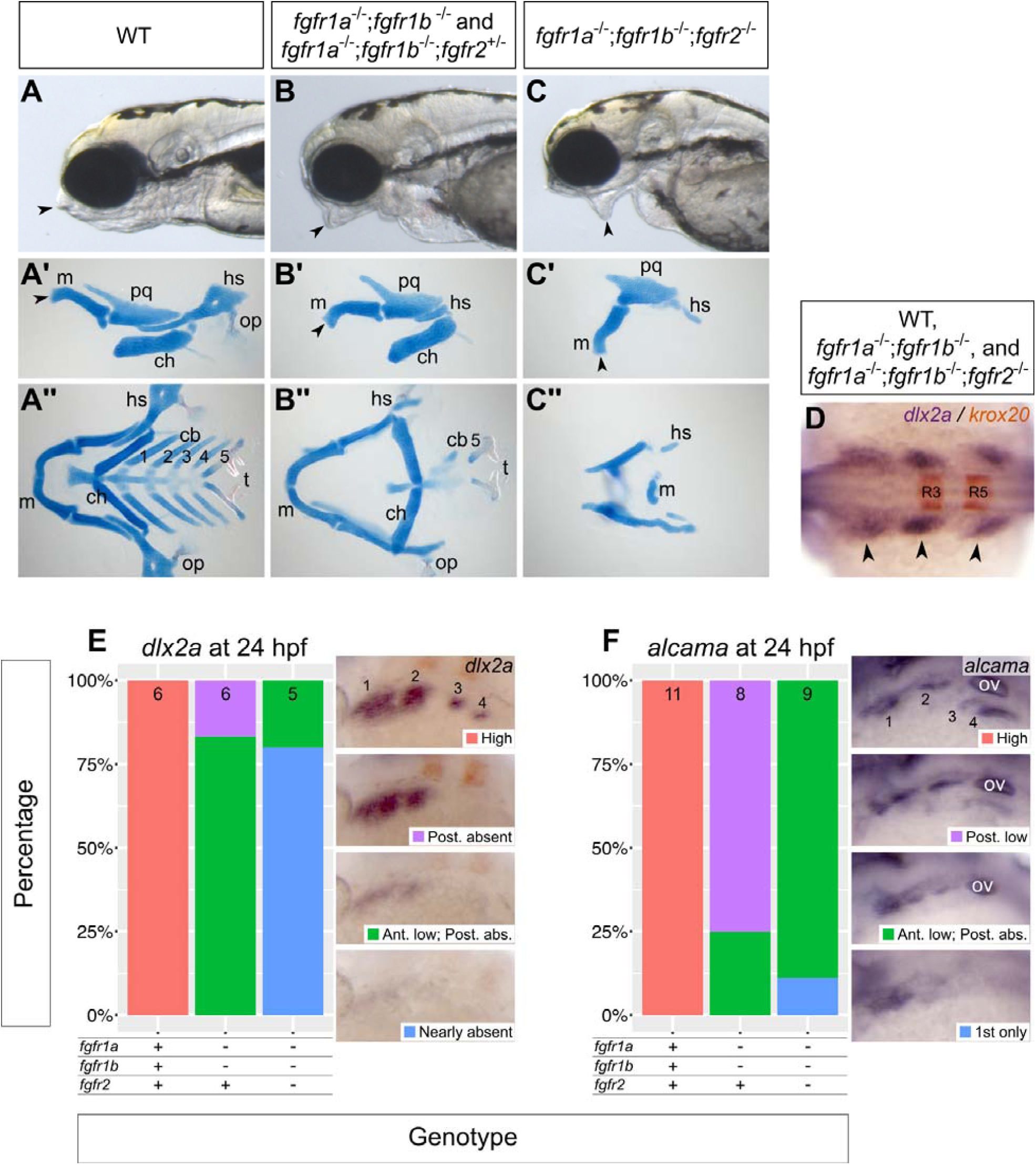
*fgfr1a, fgfr1b*, and *fgfr2* function redundantly to regulate viscerocranial development. (A, B, C) Lateral view of 5 dpf wild-type (WT, A; n = 12), *fgfr1a*^-/-^;*fgfr1b*^-/-^ (n = 3) / *fgfr1a*^-/-^;*fgfr1b*^-/-^;*fgfr2*^+/-^ (B; n = 6), *fgfr1a*^-/-^;*fgfr1b*^-/-^;*fgfr2*^-/-^ (C; n = 7) mutant larvae. (A’-C’’) Alcian blue cartilage stains of 5 dpf larvae; arrowheads noting corresponding jaw features between the live larvae in (A, B, C) and lateral view cartilage mounts in (A’, B’, C’); m = Meckel’s cartilage; pq = palatoquadrate; hs = hyosymplectic; ch = ceratohyal; op = operculum; cb 1-5 = ceratobranchials; t = teeth. (D-F) Pharyngeal arch (D, E) and pouch (F) marker analysis of *fgfr* double and triple mutant embryos at the 18-somite stage (*dlx2a* in purple/*krox20* in brown labeling rhombomeres 3 (R3) and 5 (R5), D) and 24 hpf (*dlx2a*, E; *alcama*, F). Whole mount *in situ* hybridization was performed, embryos were scored for expression, and genotypes were determined post-hoc. All embryos had indistinguishable *dlx2a* expression at the 18-somite stage (D; n = 7, 26, 6, for WT, *fgfr1a*^-/-^;*fgfr1b*^-/-^, and *fgfr1a*^-/-^;*fgfr1b*^-/-^;*fgfr2*^-/-^, respectively). In (E, F), the percentage of embryos expressing particular levels of each marker gene is represented in a stacked column chart on the left (sample size for each genotype is listed at the top of each bar), and representative images of those expression levels are shown for each marker to the right (dorsolateral views, rostral to the left and dorsal up; pharyngeal arches (E) and pouches (F) are labeled 1-4; Post. = Posterior arches/pouches; Ant. = Anterior arches/pouches; abs. = absent; in (F), ov = otic vesicle).

**Fig. 7.**
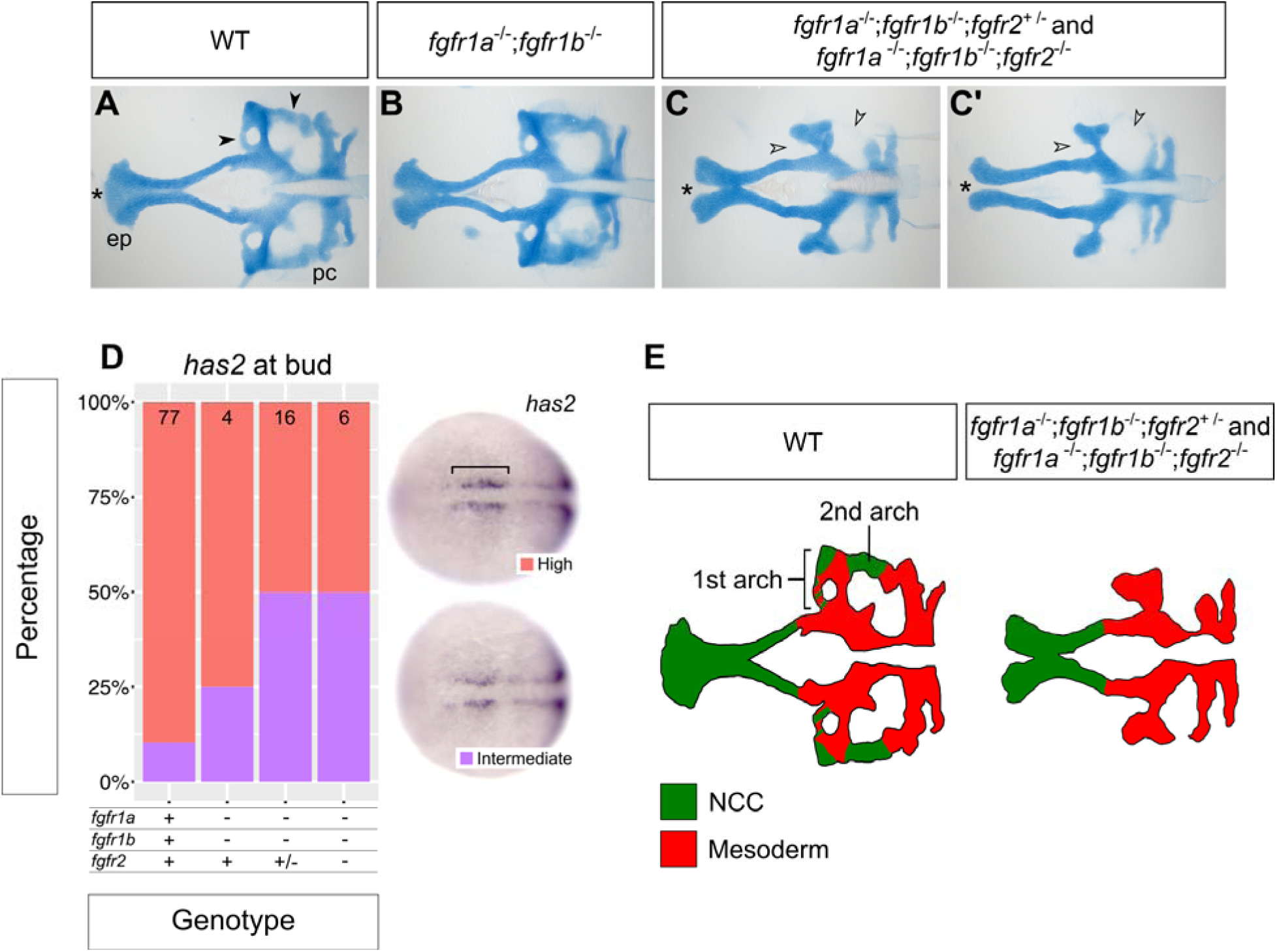
*fgfr1a, fgfr1b*, and *fgfr2* function redundantly to regulate neurocranial development. (A, B, C’) Alcian blue cartilage staining of 5 dpf wild-type (WT, A; n = 12), *fgfr1a*^-/-^;*fgfr1b*^-/-^ (B; n = 3), *fgfr1a*^-/-^;*fgfr1b*^-/-^;*fgfr2*^+/-^ (n = 6) / *fgfr1a*^-/-^;*fgfr1b*^-/-^;*fgfr2*^-/-^ (n = 7) (C, C’) mutant larvae; ep = ethmoid plate; pc = parachordal neurocranium. Notice the variable fusion of the trabeculae (*) in *fgfr1a*^-/-^;*fgfr1b*^-/-^;*fgfr2*^-/-^ triple mutants (C, C’) compared to WT (A); full fusion in 3/7 animals, partial fusion in 2/7 animals, no fusion in 2/7 animals. Open arrowheads in (C, C’) denote missing regions of the parachordal neurocranium (compare to filled arrowheads in (A)). (D) Cephalic mesoderm marker analysis of *fgfr* double and triple mutant embryos at the bud stage using *has2*. Whole mount *in situ* hybridization was performed, embryos were scored for expression, and genotypes were determined post-hoc. The percentage of embryos expressing “High” or “Medium” *has2* expression is represented in a stacked column chart on the left (sample size for each genotype is listed at the top of each bar), and representative images of those expression levels are shown for each marker to the right (dorsal views, rostral to the left; bracket denotes cells specified for cephalic development. (E) Traces of cartilage mounts in (A) and (C), filled in with expected lineage contributions, adapted with permission from McCarthy et al., 2016.

Recent studies have proposed that during development of the head skeleton, Fgf signaling is required either during pharyngeal pouch/arch formation or maintenance (Crump et al., 2004a), or later during cartilage formation (Larbuisson et al., 2013). We therefore asked at which stage our Fgfr mutations affect cranial development by assessing the expression of known marker genes of pharyngeal endoderm and NCCs, the cells that primarily form the pharyngeal pouches and arches, respectively. In a wild-type 18-somite stage embryo, NCCs expressing *dlx2a* have migrated into the pouches where they form three distinct clusters on each side of the embryo, at the anterioposterior level of the midbrain and hindbrain (arrowheads, Fig. 6D). The most anterior of these clusters will give rise to the first pharyngeal arch, the middle to the second arch, and the most posterior will later separate to give rise to arches 3-7. Interestingly, embryos of all genotypes, including *fgfr1a;fgfr1b;fgfr2* triple mutants, have normal *dlx2a* expression at the 18-somite stage, indicating that NCCs successfully migrate to and populate the pharyngeal pouches (Fig. 6D).

In a wild-type 24 hpf embryo (prim-5 stage), the most posterior cluster of NCCs has begun to separate into two distinct domains (labeled 3 and 4 in Fig. 6E), and *alcama*-labeled pharyngeal endoderm, which separates the arches, also separates into two domains posteriorly (labeled 3 and 4 in Fig. 6F). In all 24 hpf *fgfr1a;fgfr1b* double and *fgfr1a;fgfr1b;fgfr2* triple mutant embryos, however, *dlx2a* and *alcama* expression is either reduced or not detected, with the largest reduction seen in the posterior-most arches and pouches, respectively (Fig. 6E,F). Together these data suggest that NCC migration into the pharyngeal pouches occurs normally in mutants, but that these cells are not maintained. These data therefore argue for an early role of Fgf signaling in cranial development similar to that previously proposed by Crump et al. (2004).

Previous fate-mapping studies have indicated that the postchordal neurocranium is primarily derived from mesoderm, and not from NCC and that development of this tissue is dependent on Fgf8a and Fgf3 function (McCarthy et al., 2016). We therefore asked if the posterior neurocranium defects that we observe in our *fgfr1a*^-/-^*;fgfr1b*^-/-^*;fgfr2*^+/-^ and *fgfr1a*^-/-^;*fgfr1b*^-/-^;*fgfr2*^-/-^ receptor mutants are similarly due to loss of mesodermal derivatives. We used *in situ* hybridization to assess the expression of *has2*, which in wild-type bud-stage embryos is expressed in cephalic mesoderm precursor cells that localize to discrete bilateral domains flanking the anterior midline (bracket in Fig. 7D) (Camenisch et al., 2000; McCarthy et al., 2016). Unlike *fgf8a*;*fgf3* double mutants, which have reduced or no detectable expression of *has2* (McCarthy et al., 2016), we found no detectable difference in the expression of *has2* between wild-type and *fgf* receptor mutants. This result suggests that loss of Fgfr1a, Fgfr1b and Fgfr2 function does not affect the initial formation of the cephalic mesodermal, but instead is required for maintenance of this tissue, similar to what we found for the NCC during jaw development.

## Conclusions

The Fgf signaling pathway regulates numerous processes throughout development. Although specific Fgf ligands are required for many of these processes, the identity of the receptors that mediate particular cellular interactions often remain elusive. Here we have described the generation and initial characterization of mutations in each of the five zebrafish Fgf receptor genes. We showed that all single mutants are viable and fertile as adults, but that double and triple mutant combinations have defects similar to those of known ligand mutants. This has therefore allowed us for the first time in zebrafish to identify which receptors interact with select ligands. One surprise is that all phenotypes described here require loss of Fgfr1a function, suggesting that this receptor is of prime importance for Fgf-dependent processes in early development. It is interesting that Fgfr1a is also the predominant maternally supplied receptor, and as such is likely uniformly expressed in all cells during early development.

In most cases, estimating relevant receptor-ligand selectivity has relied on *in vitro* mitogenic assays, where ligands are tested for their ability to induce the proliferation of cells expressing a particular receptor or receptor isoform (Ornitz et al., 1996; Zhang et al., 2006). In only a few cases have probable receptor-ligand interactions been investigated by *in vivo* mutational analysis. Our *in vivo* analysis suggests that there is not a one-to-one relationship, where each ligand interacts with a single receptor, but that each ligand appears capable of interacting with multiple receptors. It remains to be seen if these ligands interact with only homodimers of specific receptors, or if ligand binding is able to induce heterodimerization between different receptors that are expressed in the same cell.

### Genetic redundancy or transcriptional adaptation?

We anticipated that we would find genetic redundancy between *fgfr1a* and *fgfr1b* as these ohnologs arose from a more recent genome duplication event than that gave rise to the other receptor orthologs. The apparent genetic redundancy between the *fgfr* genes was especially striking in light of previous reports that morpholino knockdown of single *fgfr* genes can result in morphological abnormalities. For example, morpholinos that block translation or splicing of *fgfr1a* were shown to cause deformation of the MHB and pharyngeal cartilages (Larbuisson et al., 2013; Scholpp et al., 2004). Likewise, morpholino knockdown of *fgfr2* was reported to cause viscerocranial cartilage and left/right asymmetry defects (Larbuisson et al., 2013; Liu et al., 2011). It is possible that the morpholino gene knockdowns do not represent the true receptor loss-of-function phenotypes. Alternatively, it is possible that transcriptional adaptation is masking more severe phenotypes in our indel mutations (Rossi et al., 2015). However, with the exception of Fgfr4 mutants, we do not detect significant changes in wild-type *fgfr* RNA expression in our *fgfr* single mutants, suggesting that the lack of phenotype in these animals is due to redundancy and not transcriptional adaptation (Fig. 2A-H). In regards to our double and triple mutant analysis, we detect significant upregulation of wild-type *fgfr3* and *fgfr4* RNA, raising the possibility that the phenotypes we describe could be less severe than those caused by alleles that do not compensate. Regardless, our results clearly identify Fgf receptors that function in specific developmental processes.

We have shown several developmental processes that use (or can use) multiple Fgf receptors. This is not unique to zebrafish as receptor redundancy has also been shown in mammals. For example, during lung development in mice, *Fgfr3*^-/-^;*Fgfr4*^-/-^ double mutants exhibit disrupted alveogenesis, whereas the lungs of single mutants are normal (Weinstein et al., 1998). Additionally, Zhao and colleagues used conditional knockout of *Fgfr1* and *Fgfr2* in concert with *Fgfr3* mutation to show that these three receptors act redundantly during mouse lens development (Zhao et al., 2008). It is possible that this type of redundancy exists elsewhere in the mouse as well. However the early embryonic lethality of *Fgfr1* and *Fgfr2* mutations makes this a difficult area of study. Going forward, the zebrafish is an attractive model to investigate these questions, in the context of Fgf signaling.

## Materials and Methods

### Husbandry

The wild-type strain *AB was used for the generation of *fgfr3*^*uc51*^ *and fgfr4*^*uc42*^. The wild-type strain NHGRI-1 was used for the generation of *fgf1a3*^*uc61*^, *fgfr1b*^*uc62*^, and *fgfr2*^*uc64*^. Zebrafish husbandry was performed as previously described (Leerberg et al., 2017; Westerfield, 2000).

### Generation of alleles

For *fgfr3*^*uc51*^ and *fgfr4*^*uc42*^, sgRNAs were designed using zifit.partners.org/ZiFiT/. Two oligonucleotides (see Table S1) were annealed and cloned into plasmid pDR274 (Addgene Plasmid #42250). For *fgfr1a*^*uc61*^, *fgfr1b*^*uc62*^, and *fgfr2*^*uc64*^, sgRNAs were designed using CRISPRscan (Moreno-Mateos et al., 2015) and produced as described (Shah et al., 2016). Briefly, overlap PCR of a T7 RNA promoter containing gene-specific oligonucleotide and a PAGE-purified scaffold oligonucleotide (see Table S1) was used to generate the DNA template for *in vitro* transcription. *cas9* mRNA was produced using the pT3TS-nls-zCas9-nls containing a codon-optimized Cas9 with two nuclear localization sequences as described in (Jao et al., 2013). The plasmid was linearized with XbaI and transcribed using the mMESSAGE mMACHINE T3 Transcription Kit (Thermo Fisher, Cat. No. AM1348). The sgRNA and *cas9* mRNA were coinjected into one-cell embryos with phenol red (5% in 2M KCl) at concentrations of 60 ng/μL and 30 ng/μL, respectively.

CRISPR efficiency was determined by comparing the targeted loci of eight injected embryos and eight uninjected control embryos (24 hpf) by High Resolution Melt Analysis (HRMA) as described (Dahlem et al., 2012) (see Table S2 for primers used). Germline mutations were identified by PCR analysis of sperm DNA from injected males (see Table S2 for primers used). Indels were sequenced, and individuals containing frameshift inducing indels were outcrossed to *AB or NHGRI-1 to obtain F1s. To reduce the potential for off-target effects, all lines were outcrossed at least four times prior to analysis.

### Genotyping

Genomic DNA was extracted from caudal fin tissue. Genotypes were determined using standard PCR conditions (*fgfr1a*^*uc61*^, *fgfr1b*^*uc62*^, *fgfr2*^*uc64*^, and *fgfr3*^*uc51*^) or HRMA (*fgfr4*^*uc42*^), and primers listed in Table S2.

### RNA *in situ* hybridization

RNA probes that detect the following genes were used: *ta* (Schulte-Merker et al., 1992); *myod* (Begemann and Ingham, 2000); *pax2a* (Krauss et al., 1991); *tbx5a* (Begemann and Ingham, 2000); *fgf24* (Draper et al., 2003); *etv4* (Munchberg et al., 1999); *fgf8a* (Reifers et al., 1998). For *has2, dlx2a, nkx2.7, alcama, fgf10a* and *en2a* probe synthesis, mRNA was isolated from 24 hpf embryos using TRI reagent (Sigma-Aldrich, Cat. No. T9424) and synthesized into cDNA using the RETROScript Reverse Transcription Kit (ThermoFisher, Cat. No. AM1710). DNA templates for *in vitro* transcription were PCR amplified with Phusion polymerase (New England BioLabs, Cat. No. M0530L) and primers found in Table 4. Reverse primers contained a T7 RNA polymerase promoter (Table S3, underlined portion), and *in vitro* transcription yielded antisense probes (Roche T7 RNA polymerase, Cat. No. 10881775001). Probes were purified with the RNA Clean and Concentrator kit (Zymo, Cat. No. R1015) and G-50 sephadex columns (GE Healthcare, Cat. No. 45-001-398). Probes were used at a concentration of 0.5-2ng/μL in hybridization solution.

Samples were fixed in 4% paraformaldehyde (PFA) overnight at 4°C or 4 hours at room temperature. Samples were dehydrated with 100% methanol and stored at - 20°C for at least 16 hours. Embryos >30 hpf were bleached for ∼10 minutes prior to proteinase K digestion in 3% H_2_O_2_, 0.5% KOH. Color *in situ* hybridizations were performed similar to Thisse and Thisse (Thisse, 2008), with the exception that 5% dextran sulfate was included in the hybridization solution.

### Bone and cartilage stains

For the scale stain depicted in Fig. S2, adults were fixed in 4% paraformaldehyde for 3 days, followed by 3 × 30 minute rinses in deionized water. Samples were bleached with 0.5% H_2_O_2_/1% KOH to remove pigment, and scales were stained with 1% alizarin red/1% KOH. Cartilage stains in Figs 5 and 6 were performed as previously described (Walker et al., 2006).

### Imaging

Embryos were mounted in 4% methylcellulose and imaged on a Leica MZ16 F stereomicroscope. Embryos in Fig. 3E-G were flat-mounted in 70% glycerol and imaged on a Zeiss Axiophot microscope. Images were collected with a Leica DFC500 digital camera.

### RT-qPCR

For RT-qPCR experiments, tail biopsies were collected from 24 hpf embryos and placed immediately in genomic DNA lysis buffer (10 mM Tris pH 8.4, 50 mM KCl, 1.5mM MgCl_2_, 0.3% Tween-20, 0.3% NP-40). Bodies were collected in 1.6 mL microcentrifuge tubes and stored in liquid nitrogen until tail tissue could be genotyped using the primers listed in Table 3. Total RNA was isolated from the bodies according to (de Jong et al., 2010) with the following exceptions: disposable pestles (USA Scientific, Cat. No. 1415-5390) were used in lieu of metal; Tri Reagent (Thermo Fisher Scientific, Cat. No. AM9738) in lieu of Qiazol; 100 μL of chloroform was added to homogenate instead of 60 μL; vacuum grease was used in lieu of phase lock gel heavy; RNA Clean and Concentrator with on column DNase treatment (Zymo Research, Cat. No. R1014) in lieu of RNeasy MinElute Cleanup; RNA was eluted with 8.5 μL nuclease-free water instead of 14 μL. RNA was quantified using a NanoDrop 1000 (Thermo Fisher Scienctific). cDNA libraries were prepared using 200 ng of total RNA in a 10 μL reaction using the RETROscript Reverse Transcription Kit (Thermo Fisher Scientific, Cat. No. AM1710), and diluted 1:5 after reverse transcription.

20 μL qPCR reactions were prepared with SsoAdvanced Universal SYBR Green Supermix (Bio-Rad), 1 μL of diluted cDNA, and primers (final concentration: 0.25 μM) listed in Table S4. Only primer sets with PCR efficiency between 1.9 and 2.1, and an R^2^ value above 0.98 were used for qPCR experiments. Reactions were performed on a Bio-Rad CFX96 machine with three technical replicates. Samples whose technical replicates had a standard deviation greater than 0.26 cycles were discarded. Fold change between wild-type and mutant animals was determined using the ddCt method and *rpl13a* as the reference gene. Unpaired Student’s T-Tests were performed to determine statistical significance.

### Statistics and plotting

Statistics were performed in R and RStudio, using standard packages (R Core Team, 2015). Graphing was performed in R, using ggplot2 (Wickham, 2009).

## Acknowledgements

We thank Kira Lin and Lan-Uyen Nguyen for assistance identifying and maintaining the alleles used in this report. We are grateful to members of the Draper, C. Erickson, P. Armstrong, L. Jao, S. Burgess, and C. Juliano labs (UC Davis) for valuable discussion of this work, and to Dr. Carol Erickson, Matt McFaul, and Sydney Wyatt for their critical review of this manuscript.

## Supplemental Material

**Fig. S1.**
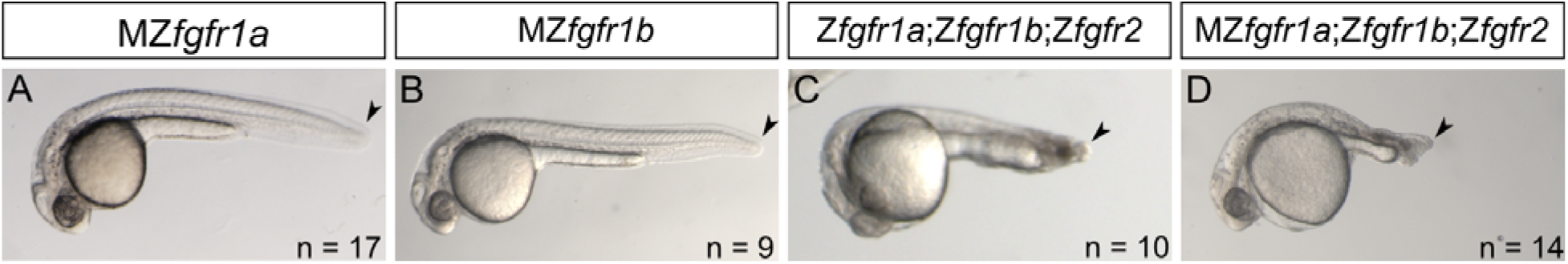
Posterior mesoderm development in MZ*fgfr1a*, MZ*fgfr1b*, and *fgfr1a*;*fgfr1b*;*fgfr2* triple mutants is normal. (A, B) 30 hpf embryos missing both maternal and zygotic contributions of *fgfr1a* (A) or *fgfr1b* (B) exhibit normal posterior mesoderm development. (C) 24 hpf embryo missing zygotic contributions of *fgfr1a, fgfr1b* and *fgfr2*. Compare to Fig. 3B. (D) 28 hpf embryo missing maternal and zygotic contributions of *fgfr1a* and zygotic contributions of *fgfr1b* and *fgfr2*. Compare to Fig. 3D. Lateral views; anterior is to the left, dorsal is up.

**Fig. S2.**
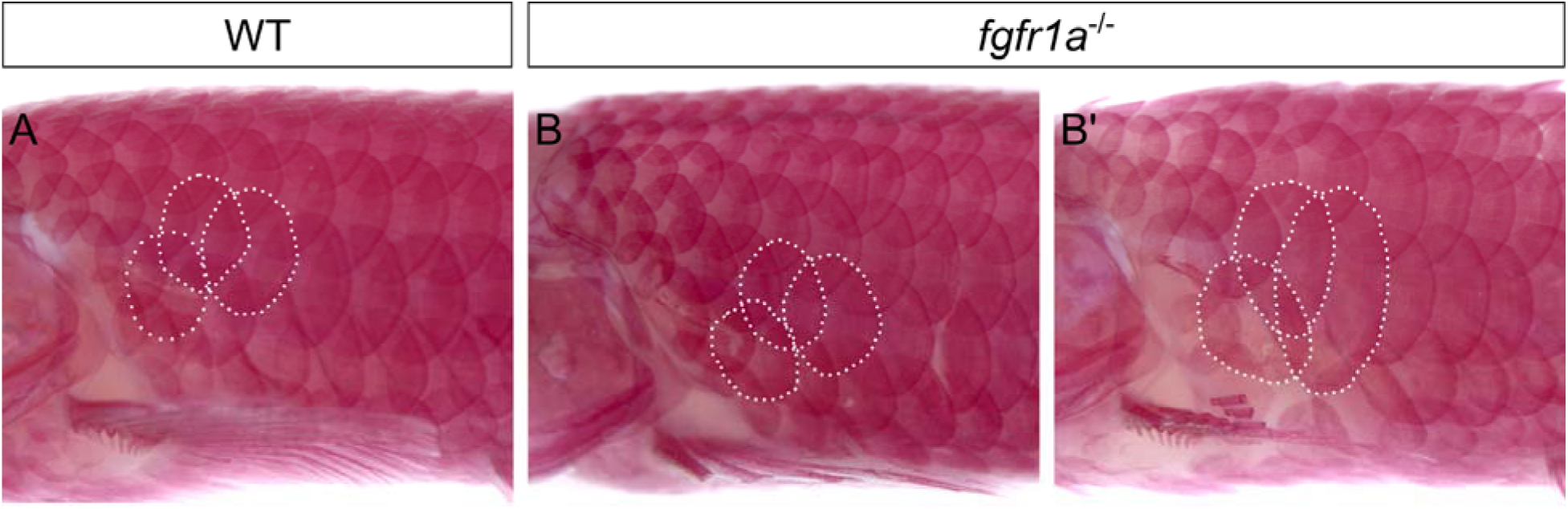
*fgfr1a*^uc61^ mutants have a variable and subtle defect in scale development. (A-B’) Wild-type (A) and *fgfr1a*^uc61^/^uc61^ (B, B’) adults stained with alizarin red. Three similarly located scales are outlined in white in each panel. In 2/5 *fgfr1a*^uc61^ mutants analyzed, scales appeared normal in size (B). In 3/5, however, select flank scales were larger (B’). Anterior is to the left, dorsal is up.

**Table S1.**
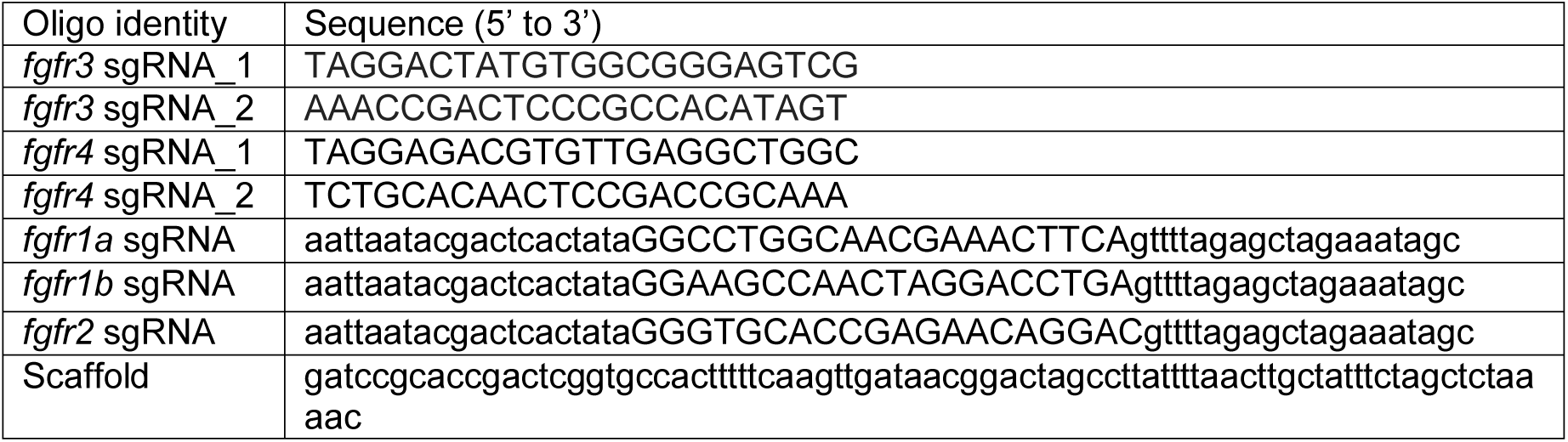
Oligonucleotide used for sgRNA construction.

**Table S2.**
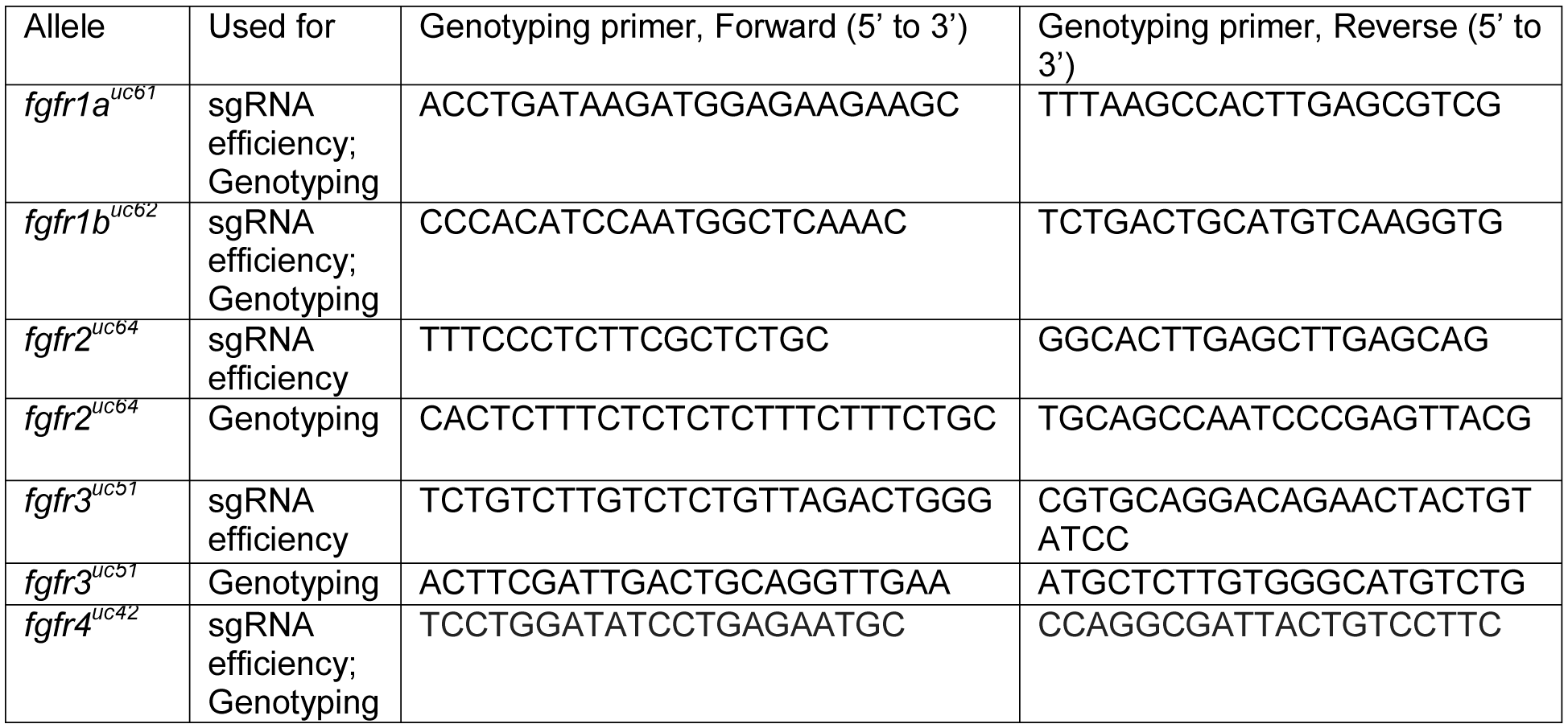
Genotyping primers.

**Table S3.**
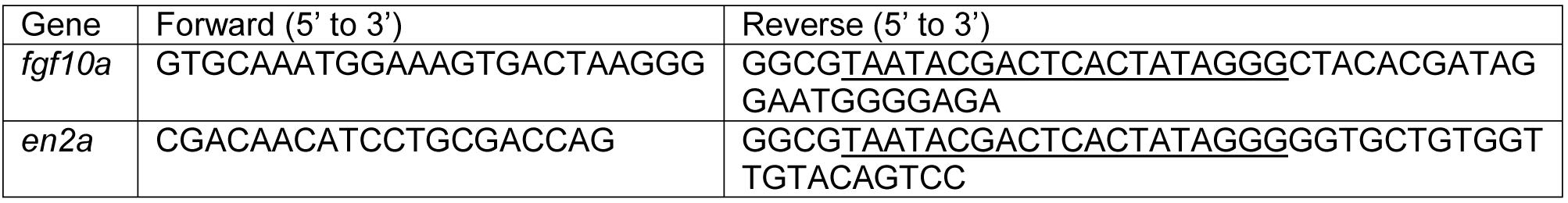
Primers used for RNA *in situ* hybridization probe synthesis

**Table S4.**
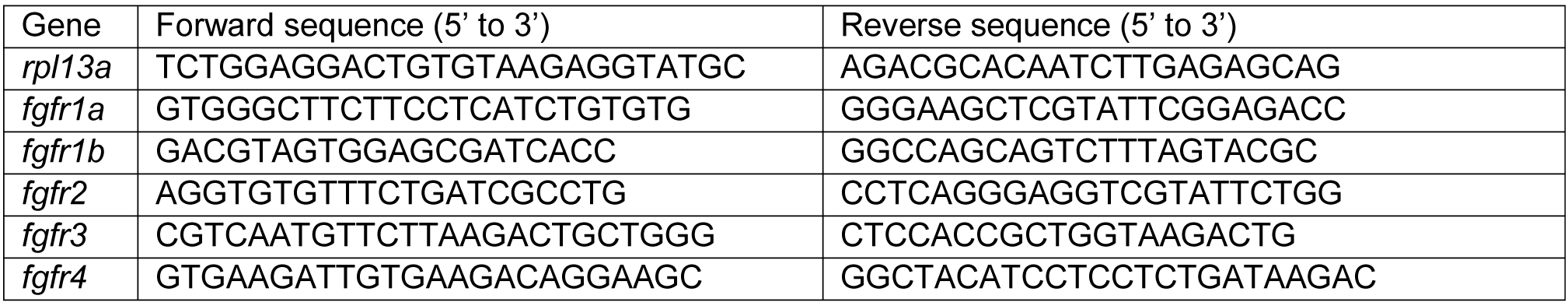
qPCR primers.

